# Rapid T cell receptor interaction grouping with ting

**DOI:** 10.1101/2020.05.04.069914

**Authors:** Felix Mölder, Ulrik Stervbo, Lucie Loyal, Petra Bacher, Nina Babel, Sven Rahmann

**Affiliations:** Genome Informatics, Institute of Human Genetics, University of Duisburg-Essen, 45147 Essen, Germany; Institute of Pathology, University of Duisburg-Essen, 45147, Germany; Center for Translational Medicine, Medical Clinic I, Marien Hospital Herne, Ruhr-University Bochum, 44623 Herne, Germany; Si-M / “Der Simulierte Mensch” a science framework of Technische Universität Berlin and Charité - Universitätsmedizin Berlin, Berlin, Germany; Regenerative Immunology and Aging, BIH Center for Regenerative Therapies, Charité Universitätsmedizin Berlin, Berlin, Germany; Institute of Immunology, Christian-Albrechts Universität zu Kiel and Universitätsklinik Schleswig-Holstein, Kiel, Germany; Institute of Clinical Molecular Biology, Christian-Albrechts Universität zu Kiel, Kiel, Germany; Charité – Universitätsmedizin Berlin, Corporate Member of Freie Universität Berlin, Humboldt-Universität zu Berlin; Berlin Institute of Health, Berlin-Brandenburg Center for Regenerative Therapies, Augustenburger Platz 1, 13353 Berlin, Germany

## Abstract

**Motivation:** Clustering of antigen-specific T cell receptor repertoire (TCRR) sequences is challenging. The recently published tool GLIPH aims to solve this problem. However, clustering large repertoires takes several days to weeks, making its use impractical in larger studies. In addition, the methodology used in GLIPH suffers from several shortcomings, including non-determinism, potential loss of significant antigen-specific sequences or inclusion of too many unspecific sequences.

**Results:** We present an algorithm for clustering TCRR sequences that scales efficiently to large repertoires. We clustered 26 real datasets with up to 62 000 unique CDR3*β* sequences using both GLIPH and an implementation of our method called ting. While GLIPH required multiple weeks, ting only needed about one hour for the same task. In addition, we found that in naïve repertoires, where no or very few antigen-specific CDR3 sequences or clusters should exist, our method indeed selects fewer sequences.

**Availability:** Our method has been implemented in Python as a tool called ting, using numpy and NetworkX. It is available on GitHub (https://github.com/FelixMoelder/ting) and on PyPI under the MIT license.

## Introduction

T cells are an essential part of the immune system of higher vertebrates. Each T cell is endowed with its own particular heterodimeric T cell receptor (TCR), which conveys the ability to bind to distinct antigenic peptides presented in the context of the major histocompatibility complex. The hypervariable complementary-determining region 3 (CDR3) of both TCR subunits is responsible for the antigen specificity (1,2). Understanding the relationship between the TCR CDR3 sequence and the antigen could have a major impact on diagnosis and tailor-made cancer immunotherapies. This has, however, proven challenging due to the plasticity and crossreactivity of the TCRs (2). Recently, Glanville et al. (1) proposed the method GLIPH that clusters TCR sequences sharing the same antigen-specificity by global and local sequence similarity.

When applying GLIPH to several thousand TCR*β* sequences, we observed that its method for identifying significantly enriched *k*-mers is a runtime intensive and non-deterministic process yielding slightly varying *k*-mers in every run. Additionally, the local clustering step could take several days or weeks to complete. We therefore devised an improved algorithm for clustering TCR sequences by local similarity and a more robust, deterministic method for the detection of significantly enriched motifs. The improvement presented here allows us to reduce the run time significantly from weeks to just a few hours. We further found that GLIPH contained an error that may incorrectly overlook some significantly enriched *k*-mers, in particular those that do not appear at all in a control set. Moreover, evaluation of the resulting clusters showed unexpectedly large cluster sizes on naïve TCR repertoires (TCRRs). The method presented here produces clusterings that conform better to expected cluster sizes on naïve TCRRs.

## Methods

The input to the TCR repertoire (TCRR) clustering problem consists of a a set *S* of sample amino acid sequences (for example CDR3*β* regions) and a control set *N* of corresponding amino acid sequences from a naïve repertoire (or several ones). The desired *output* is a partitioning of *S*, i.e. a set *C* = {*C*_1_,*C*_2_,…,*C_z_* } such that each *cluster C_i_* is a subset of *S*, the *C_i_* are pairwise disjoint (*C_i_* ⋂ *C_j_* = {} for all *i* ≠ *j*), and their union forms the whole set 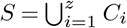.

As described by Glanville et al. (1), one expects CDR3 sequences to be joined in the same cluster that are specific to the same antigen. So a naïve repertoire should mostly consist of very small unspecific clusters, even singletons, while the repertoire after an activated immune response should contain larger antigen-specific clusters. Before we present our algorithm, we give an overview of GLIPH’s approach (1).

### Summary of the GLIPH approach

First, the number of occurrences of each peptide of length 2, 3 and 4, referred to as *motif,* is counted separately in the sample set *S* and in the control set *N*. A randomized resampling-based criterion is used to extract motifs that are significantly overrepresented in *S* compared to *N*. This leads to a set *M*\* of significant motifs. A *local clustering* of the sequences in *S* is performed by placing any two sequences that have a common motif in *M*^*^ into the same cluster. Furthermore, a *global clustering* step is performed by placing any two sequences *s, s*’ from *S* in the same cluster if they have a small Hamming distance (threshold 2 if |*S*| ≤ 125, threshold 1 if |*S*| > 125). The result is the desired output: a partitioning of the CDR3*β* sequences from *S* into disjoint clusters.

In the following, we describe how each step of this approach can be improved upon methodologically and executed more efficiently.

### Finding significant motifs

The approach in GLIPH is based on resampling and empirical p-values. First, the exact counts for the motifs *M_S_* (peptides of length 2, 3 and 4 contained in sequences in *S*) are obtained. This results in a “counter” function *C_S_*: *M_S_* → ℕ. It is assumed that the naïve control repertoire *N* contains more sequences than *S*. Then, for *n* rounds, |*S*| sequences are randomly sampled from *N*, resulting in a sample set *R_j_,j* = 1,…,*n*, and the motifs *M_j_* in *R_j_* are counted, resulting in a counter function *C_j_*: *M_j_* → ℕ. For each motif *m* ∈ *M_S_*, it is determined in which fraction of samples the motif appears at least as frequently as in *S*: We call 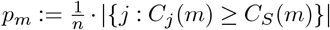 the *empiricalp-value* of motif *m* (after *n* resampling rounds). It is valid to compare the counts *C_j_* (*m*) and *C_S_*(*m*) because *R_j_* and *S* are of the same size |*S*|. A motif *m* is then *significantly enriched* if all of the following conditions are true:

1. The motif is seen at least 3 times in *S*, i.e., *C_S_* (*m*) ≥ 3.
2. The observed-vs-expected fold change *ρ_m_* of *m* reaches a threshold *T*, where *ρ_m_* is defined as the ratio between *C_S_*(*m*) and the average count in the naïve resamplings:

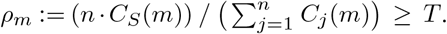 In GLIPH, *T* = 10 is used, unless *C_S_*(*m*) = 3, where *T* =100.
3. The empirical p-value is small, *p_m_* ≤ 0.001.

This methodology has the following disadvantages: First, the approach only works for |*N*| ≥ |*S*| (which is often satisfied, but there is no guarantee) because each re-sampling round must produce a down-sampled version of *N* of size |*S*|. Second, the results depend on the number *n* of re-sampling steps (*n* = 1000 is used in GLIPH). Since a large number of rounds is required for accurate estimates of *p_m_* and *ρ_m_*, this approach is slow. Third, for any finite value of *n*, the resulting set of significantly enriched motifs is non-deterministic and hence may vary with each execution. In addition, the published version of GLIPH contains an error that skips each motif *m* with 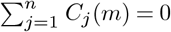, even if it satisfies all conditions, apparently to circumvent a division by zero.

We therefore suggest two alternative methods of finding significantly enriched motifs: First, we only correct the erroneous behavior but leave the method otherwise unchanged; this is referred to as the *bug-fixed* method in the Results section. This version inherits the other issues of the original algorithm: the requirement that |*N*| ≥ |*S*|, slow speed due to many resampling rounds, and non-deterministic behavior. Therefore, we secondly present an entirely different test strategy using Fisher’s exact test (3). Consider the contingency table shown in Table 1 that relates the occurrence count of motif *m* in the sets *S* and *N* to the sizes of these sets. Fisher’s exact test accommodates different sizes |*S*| and |*N*| without constraints or resampling and deterministically yields a p-value for the null hypothesis that the rows (or columns) of the contingency table are independent, which corresponds to the statement that 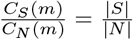 or equivalently 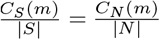.

**Table 1.**
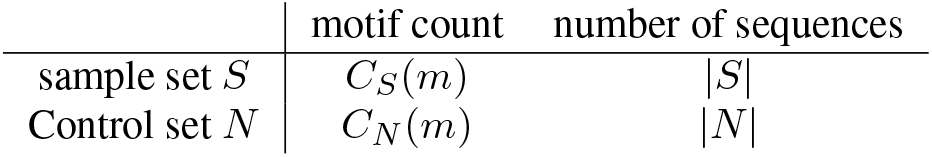
Contingency table for discovering significantly enriched motifs *m* with Fisher’s exact test.

We call a motif significantly enriched if the following two conditions are satisfied:

1. The fold change 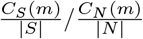 exceeds a given threshold *T* (by default set to *T* = 10)
2. The Fisher p-value falls below a given Bonferroni-corrected threshold 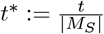, where *t* is the uncorrected p-value threshold (typically *t*:= 0.05). (The correction is conservative and accounts for the total number of tested motifs, resulting in a family-wise error rate of 5%.)

In contrast to the GLIPH criteria, we do not need a requirement on the absolute count *C_S_* (*m*) because motifs with low counts are insignificant by Fisher’s test. The two remaining criteria cover effect size (fold change) and statistical significance. In the following, *M*^*^ denotes the set of significantly enriched motifs.

### Local clustering

The goal of the clustering steps is to build a graph *G* = (*S,E*), where the nodes are the CDR3 sequences in the condition-specific repertoire *S* and the presence of an edge {*s,’*} ∈ *E* denotes global or local similarity between *s* ∈ *S* and *s’* ∈ *S*.

The local clustering step consists of placing sequences from *S* that share a motif from *M*^*^ in the same cluster, i.e., creating edges {*s,s*’} if *s* and *s’* have a common substring in *M*^*^. The approach taken by GLIPH uses an inefficient algorithm to do this: It compares every pair of sequences, scanning each pair for each motif, leading to a running time of *O*(|*M*| · |*S*|^2^), where typically |*S*| » |*M*|, yielding times of days up to weeks for a single dataset.

We improve upon this as follows.

1. We avoid redundancy among the motifs (recall that they can be of different lengths, 2, 3, or 4) by selecting short representatives.
2. We use a faster method to join sequences with common motifs.

The first step creates a directed acyclic motif substring graph *G*\* = (*M*,E*\*): An edge (*m,m*’) exists in *G*\* if and only if *m* is a substring of *m’*. We observe that only the motifs *m* that have no incoming edge need to be further considered: If there is an edge *m* → *m’* and *m’* occurs in a sequence *s* ∈ *S*, then *m* also occurs as a substring. In this way, *M*\* is reduced. The second step consists of joining sequence sets with common motifs (from the reduced *M*\*). This is done using a disjoint-set (or union-find) data structure (4) that maintains a partition of *S*. Clusters are implicitly defined by assigning a representative sequence to each cluster. If *S* = {*s*ι,…,*s*?_*s*_?}, we let *r_i_* be the index of the representative sequence of the cluster containing *s_i_*. Initially, each sequence *s ∈ S* is in a singleton cluster, so *r_i_ = i* for all *i* = 1,…, |*S*|. We iterate over all motifs *m ∈ M* * and consider the sets *I_m_*:= {*i*: *m* is a substring of *s_i_*}. We set the representatives of all *i ∈ I_m_* to the representative of min(*I_m_*); thereby joining the clusters containing a sequence with substring *m* into a single cluster. Details about the union-find data structure and the implementation of its operations can be found in the textbook by Cormen et al. (5).

A final pass over the representatives (following each path *i → r_i_* → *r_ri_* → … up to its root where *r_i_* = *i*) yields the clusters.

### Running time

The initial motif clustering step takes time *O*(|*M*_2_| · |*M*_3_| · |*M*_4_|), and the sequence clustering step takes time *O*(|*M*||*S*|*α*(|*S*|)), where *α(o)* denotes the inverse Ackermann function and is a small constant in practice.

### Global clustering

We did not change the global clustering algorithm (pairwise Hamming distance computation; selection of pairs with distance below a threshold), but instead make use of vectorized functions in Python’s numpy library (6).

## Results

We compare several methods on different samples in terms of the number of significantly enriched motifs and running times for motif identification and clustering. As the number of identified motifs may differ between different methods, we measure motif identification and clustering steps separately and benchmark the clustering time using a fixed motif set. The evaluation was performed on a local workstation with a quadcore Intel(R) i7-3770 CPU at 3.40GHz and 16GB RAM.

### Compared Methods

We compare the following methods:

1. **GLIPH-original**: the original GLIPH implementation
2. **GLIPH-bugfixed**: the bug-fixed version (without any other improvements)
3. **ting-gliph**: the bug-fixed version with our fast local clustering
4. **ting-fisher**: using Fisher’s exact test with fast local clustering

We observed that GLIPH omits input sequences that do not begin with cystein. This may unintentionally remove many sequences in some datasets. For our comparison, we patched GLIPH with an option to include these sequences, and added an option to ting allowing to filter CDR3*β* sequences starting with cysteine and ending with penhylalanine, as defined by IGMT^1^. To achieve comparable results, filtering was completely turned off during the comparison.

### Samples and controls

For validating motif identification and measuring running time, we used three sets of samples. The first set holds 14 (antigen-)specific T cell samples, each containing between 673 and 2 836 unique CDR3*β* sequences. The second set consists of six large naïve samples (31 529 to 62446 unique CDR3*β* sequences), while the third set contains six samples of unknown specificity with 5 778 to 38 574 sequences.

For each sample, the repertoire of CDR3*β* sequences, was created using IMSEQ (7). As control set *N*, we used a set of naive sequences provided by GLIPH that contains 314 863 sequences^2^.

### Running times

We separately compare the times required for motif discovery (significant *k*-mers) and clustering. Figure 1 shows the running time of ting’s and GLIPH’s algorithms for motif identification in minutes. As expected, GLIPH’s original and big-fixed implementation mostly require the same time for samples, with small stochastic differences due to the non-deterministic nature of the sampling method. For a few samples, the bug-fixed implementation is observably slower because more significant motifs are discovered.

**Fig. 1.**
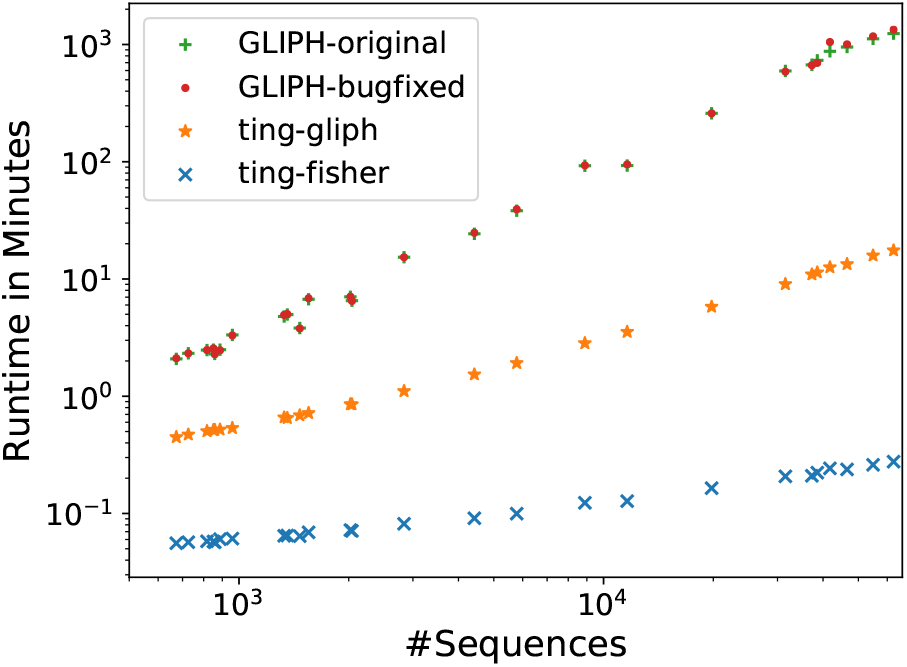
Time for identifying significant motifs applying GLIPH’s and ting’s algorithms on data sets of different sizes. Note that both axes are logarithmically scaled.

The ting implementation of GLIPH’s algorithm is faster by a factor of up to 75 ×. While GLIPH is able to process smaller samples within a few minutes, it already takes more than 40 minutes for samples with more than 6 000 sequences and up to about 22 hours for the biggest sample. In comparison, ting is able to perform motif identification for the biggest sample of 62446 sequences in about 23 minutes. In addition, we find that the deterministic approach using Fisher’s exact test is even faster. It is able to identify significant *k*-mers in less than 10 seconds even for the biggest sample. In addition, while the runtime of GLIPH’s algorithm may vary for every run due to its random resampling, the test-based algorithm has consistent running times.

Figure 2 shows the clustering runtimes required by GLIPH and ting for the different samples. As both tools used the same motif sets for clustering the calculated clusters are identical. Since clustering applying GLIPH is time consuming we were not able to obtain results for samples with more than 20000 sequences. Even for 19 789 sequences GLIPH required 6.3 days, taking more than 1000 times longer than ting. In comparison, ting is able to obtain the results in a few seconds for smaller samples and in below 2 hours for the largest one.

**Fig. 2.**
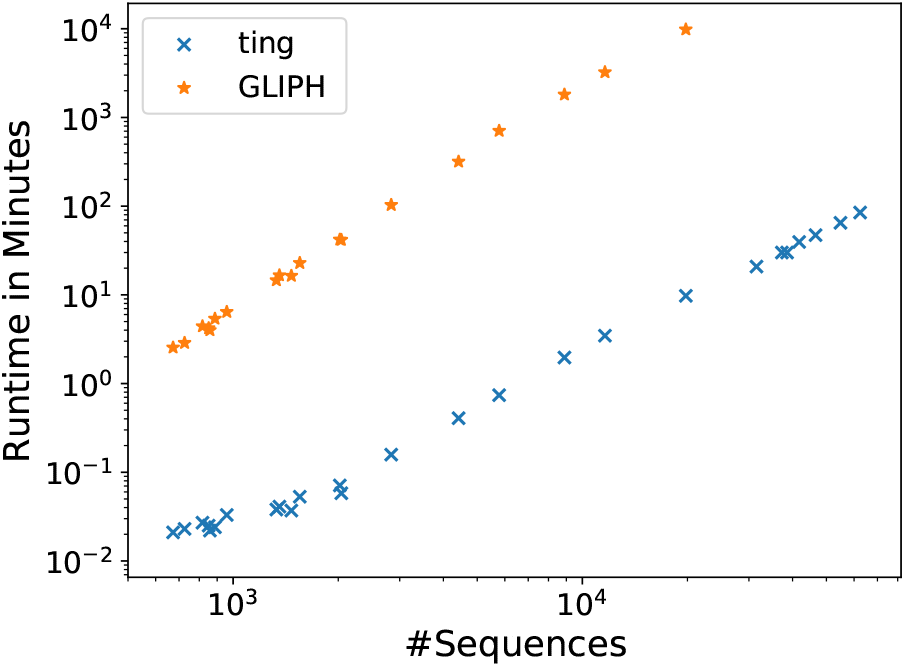
Time for clustering using GLIPH and ting on data sets of different sizes. Note that both axes are logarithmically scaled.

Even though we focused on improving the local clustering we included global clustering, as it is not possible to run the local clustering without performing global clustering steps by GLIPH.

### Identified enriched *k*-mer motifs

In order to investigate how the three motif identification algorithms differ, we compared the number of identified significant motifs from GLIPH-original, GLIPH-bugfixed (same results as tinggliph) and ting-fisher. Figure 3 compares the number of identified motifs in all samples for each pair of algorithms. Antigen-specific samples are represented by dots, naïve samples by diamonds (we should expect almost no significant *k*-mers), and samples of unknown specificity are represented by stars.

**Fig. 3.**
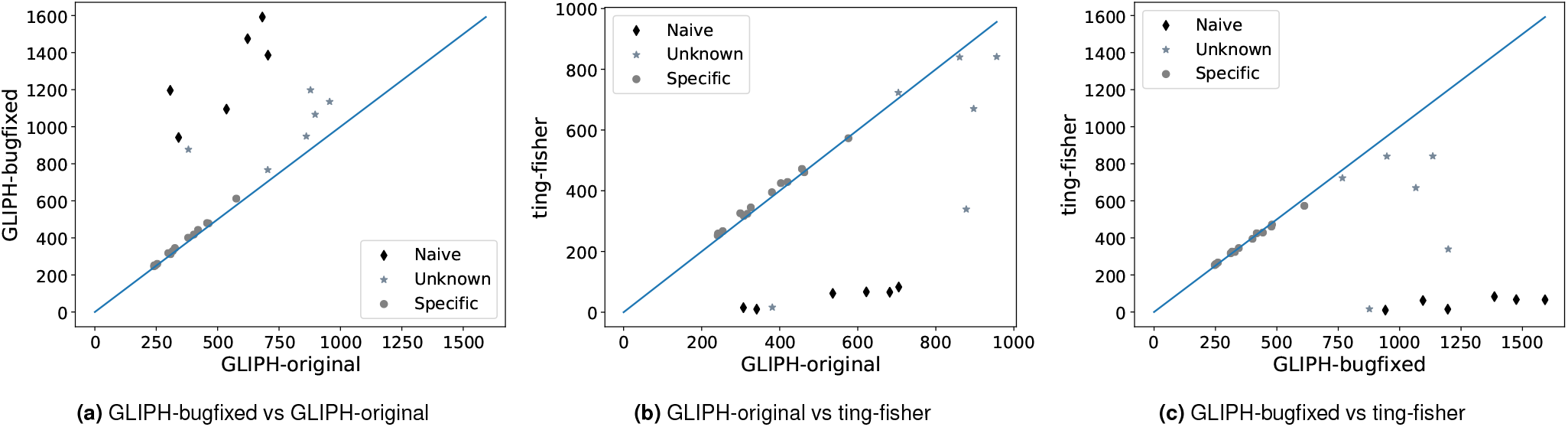
Number of significant *k*-mers identified by ting vs. GLIPH

Figure 3a shows that the number of motifs in specific samples, identified by GLIPH’s original and bug-fixed algorithm strongly correlates and that there is almost no difference. However, there is a strong difference in the number of identified motifs in the naïve samples showing that the bug-fixed algorithm identifies much more significant motifs than the original implementation. This can be explained by the nature of the bug occurring in GLIPH omitting all motifs that exist in the input sample set but not (at all) in the control set.

Comparing the results of ting-fisher to GLIPH-bugfixed (which is equivalent to ting-gliph, Figure 3b shows that naïve and specific samples are separated into two classes. While ting classifies the same amount of *k*-mer motifs as significant as GLIPH in the specific samples, the number of significant motifs identified in the naïve samples is much lower than the amount identified by GLIPH. This appears reasonable as we would expect only very few specific *k*-mers in naive samples by the definition of naive. Figure 3c shows that the ratio of identified *k*-mers between ting and the bug-fixed version of GLIPH only gets worse, where GLIPH identifies up to 7 × more significant *k*-mers in naive sample sets.

### Cluster sizes

Figure 4 shows the size of the largest cluster in relation to the whole number of sequences in each sample after using different clustering methods: Clusters were calculated separately by local clustering based on the *k*-mers identified by Fisher’s exact test and GLIPH’s algorithm, global clustering and a combination of global and local clustering using Fisher’s *k*-mers. Naive, unknown and specific samples were separately colored.

**Fig. 4.**
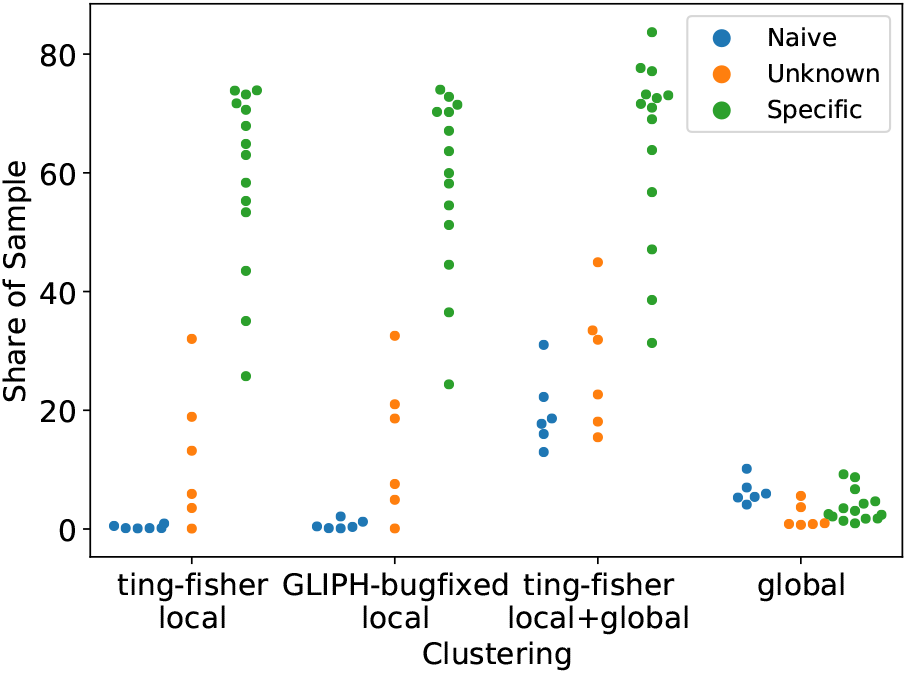
Share of largest cluster in each sample, in per cent, for different local/global clustering algorithms. Clustering was performed, from left to right, locally by Fisher’s exact test, locally by GLIPH’s bug-fixed implementation, by a combination of global and local clustering using Fisher’s exact test, and by global clustering only.

After local clustering using GLIPH or Fisher’s exact test, the size of the largest cluster is comparable. Also, the largest clusters in naive samples are relatively small, taking 2% of the whole sample at most. In contrast, the largest cluster in antigen-specific samples ranges from more than 20% to over 90%. Largest cluster sizes in samples with unknown specificity range between the sizes of naive and specific samples. Figure 3 shows that ting-fisher identifies fewer significant *k*-mers in naive samples than GLIPH-bugfixed. Therefore, one would expect that local naive clusters created by ting-fisher are smaller than clusters identified by GLIPH-bugfixed; yet this is not visible in Figure 4, which shows only the size of the *largest* cluster of each sample. Considering all clusters, GLIPH-bugfixed identifies a higher number of large clusters than ting (see supplementary results^3^). Thus cluster sizes found by toolname-fisher are in better agreement with expectations than cluster sizes identified by GLIPH-bugfixed.

Looking at the largest cluster created by global clustering draws a different picture. While global clustering builds smaller clusters than local clustering within specific samples, the clusters in naive samples are much larger. Combining local and global clustering shows, that while cluster sizes in specific samples are not much bigger compared to clusters created by local clustering only, clustsers in naive samples increase size and take up to 40% of the sample.

This leads to the impression that global clustering is not able to distinguish between naive and specific sequences and tends to cluster false positive sequences not sharing any specificity.

Even if the global clustering criteria cluster some sequences with shared specificity, it does not mean that it finds many connections that have been missed by the local criteria. It instead appears to mostly impact the false positive rate in naive samples. For this reason it seems preferable to use local clustering only.

## Conclusion

We presented ting, a similarity clustering tool for CDR3*β* sequences that implements the GLIPH method of Glanville et al., but is faster by orders of magnitude, reducing running times from potentially weeks to a few hours. This improvement enables users to analyse large datasets of tens to hundreds of thousands of sequences in reasonable time on a local PC without the need of a compute cluster, enabling many smaller labs to analyse large CDR3*β* datasets. Additionally, ting comes with a robust deterministic algorithm for identifying significant enriched *k*-mers based on Fisher’s exact test as default. Comparing local and global clustering criteria on antigen-specific and naive repertoires suggests that global clustering is unnecessary for antigen-specific samples and harmful for naive samples.

Our implementation “ting” is available on PyPI and at https://github.com/FelixMoelder/ting.

## Funding

We acknowledge funding from the EFRE.NRW program OsteoSys (EFRE-0800427; LS-1-1-019c).

## Footnotes

1 http://www.imgt.org/FAQ/#question34

2 https://github.com/immunoengineer/gliph/raw/master/gliph-1.0.tgz

3 cluster_sizes.tar.gz

## Notes

### Competing Interest Statement

The authors have declared no competing interest.

