## Supplementary figures and images for "Rapid T cell receptor interaction grouping with ting"

### naive_LULO-TC-DN1_Tc_N_CD40L-_cluster.pdf

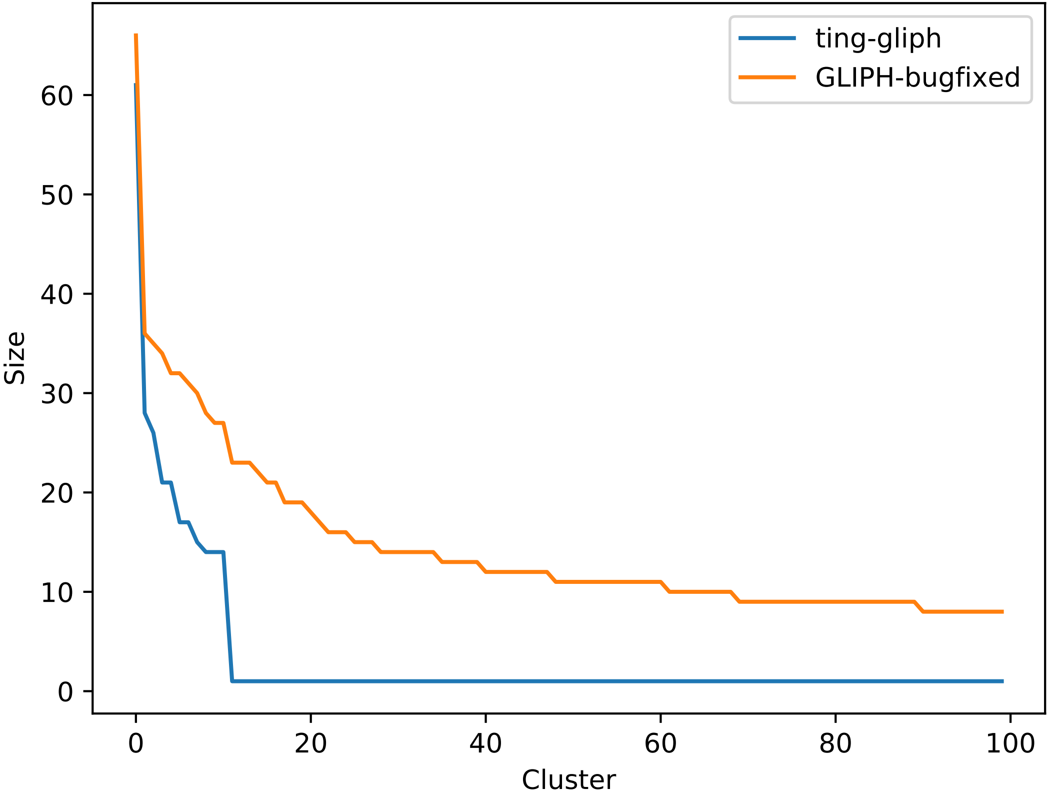

### naive_LULO-TC-DN2_Tc_N_CD40L-_cluster.pdf

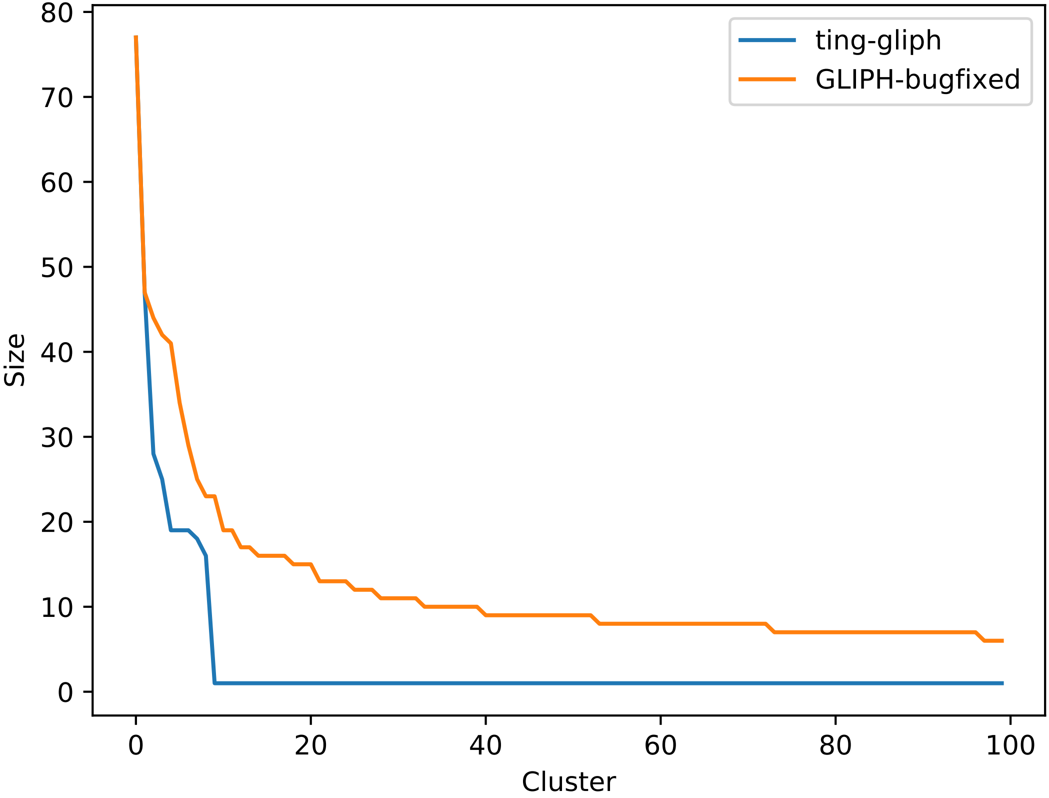

### naive_LULO-TC-DN3_Tc_N_CD40L-_cluster.pdf

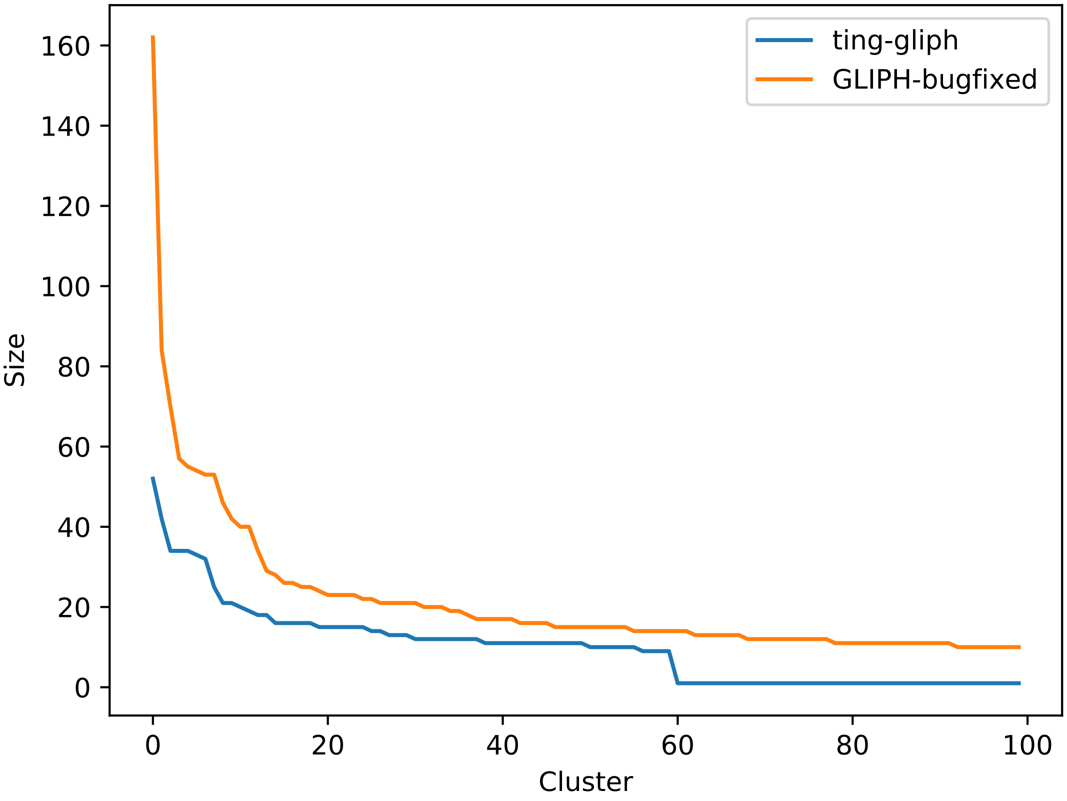

### naive_LULO-TC-DN4_Tc_N_CD40L-_cluster.pdf

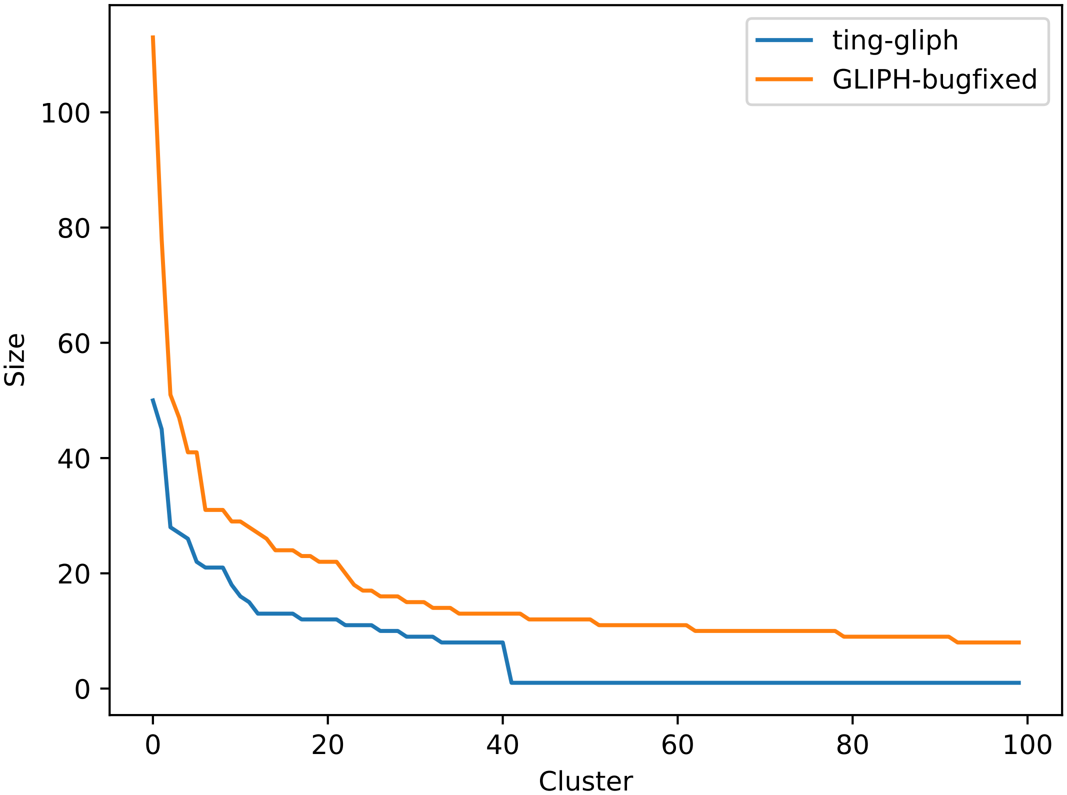

### naive_LULO-TC-DN5_Tc_N_CD40L-_cluster.pdf

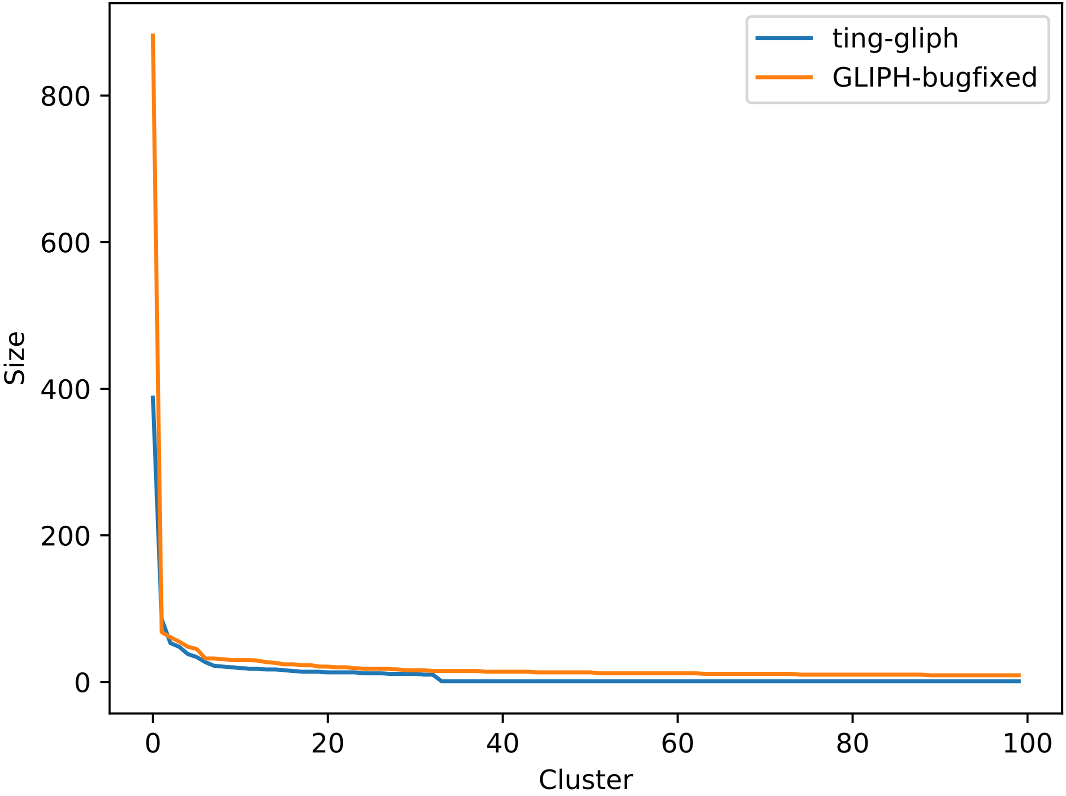

### naive_LULO-TC-DN6_Tc_N_CD40L-_cluster.pdf

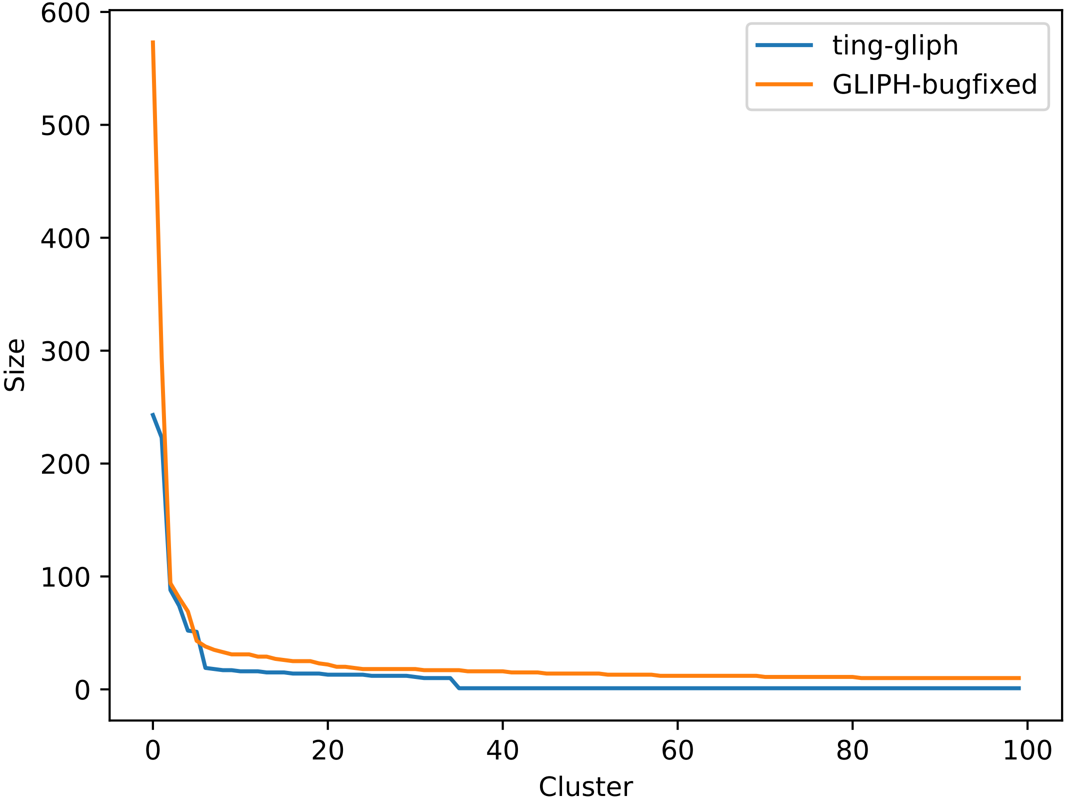

### specific_R205-L01-D704D504_cluster.pdf

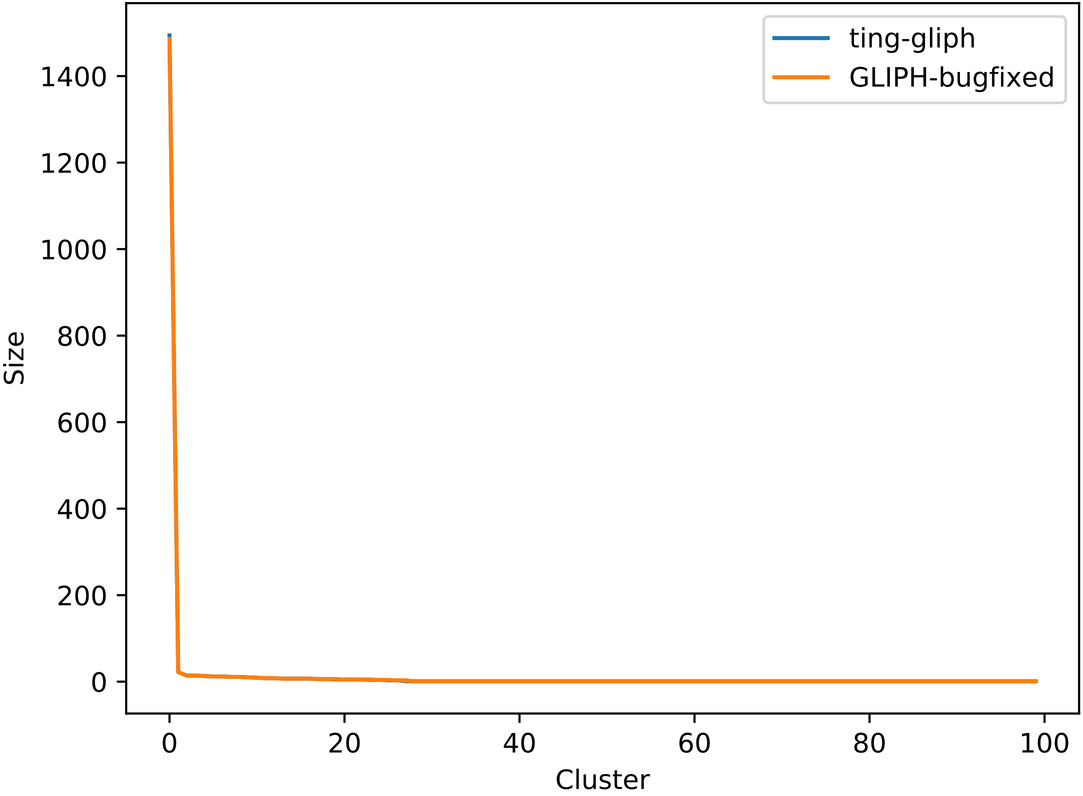

### specific_R205-L01-D706D506_cluster.pdf

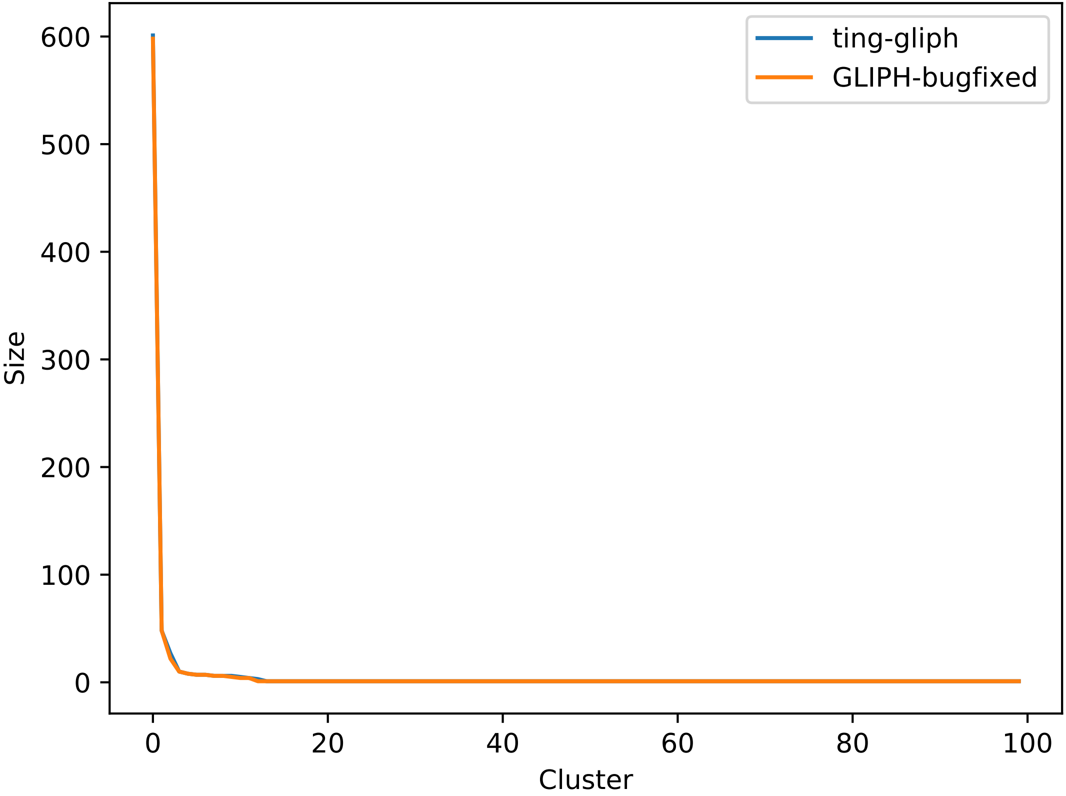

### specific_R205-L01-D709D509_cluster.pdf

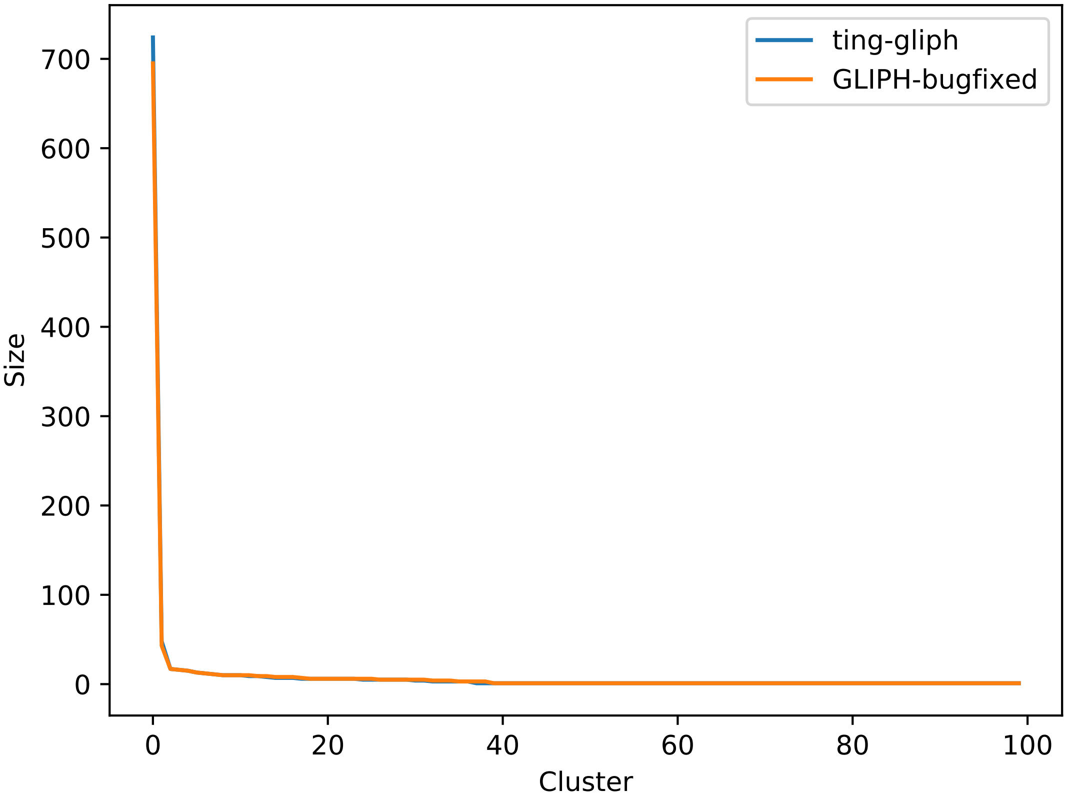

### specific_R205-L01-D712D512_cluster.pdf

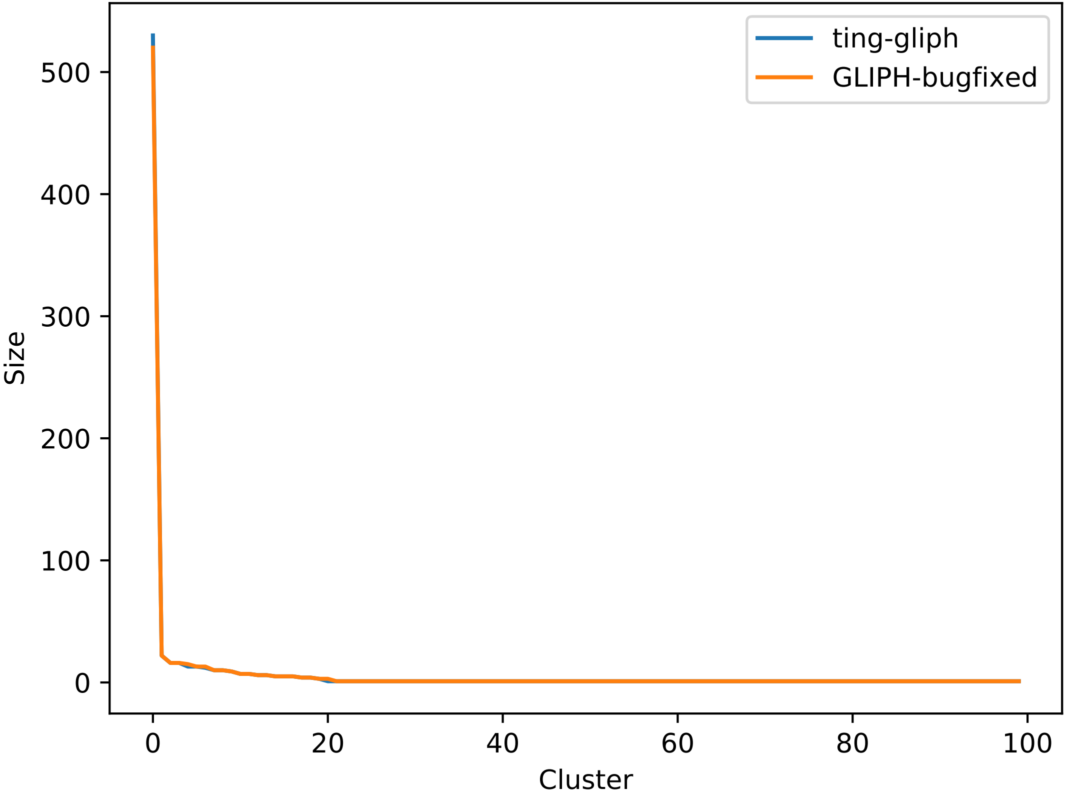

### specific_R205-L01-D715D515_cluster.pdf

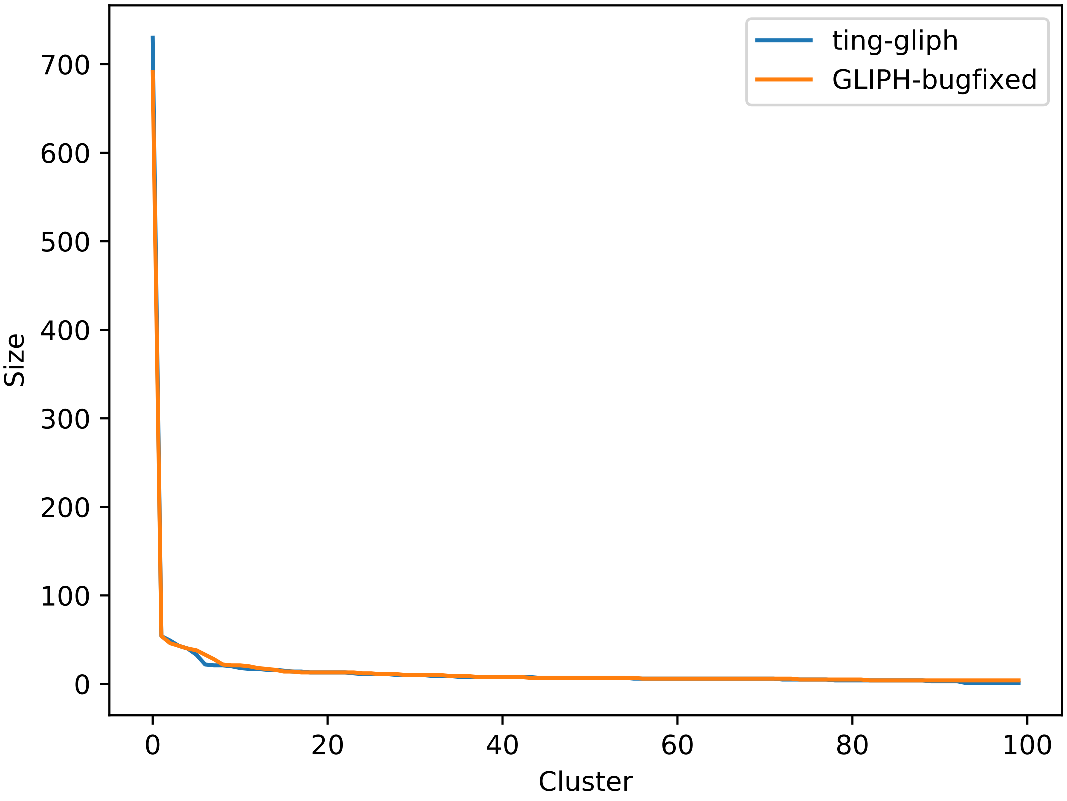

### specific_R205-L01-D716D516_cluster.pdf

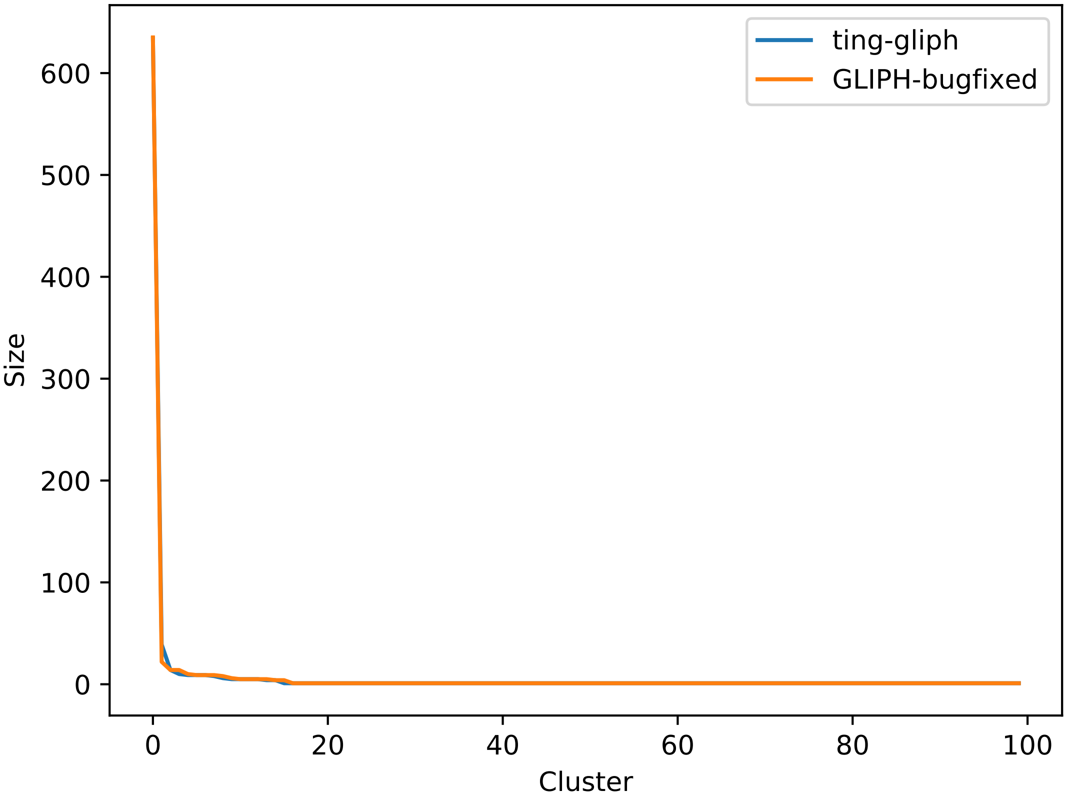

### specific_R205-L01-D727D527_cluster.pdf

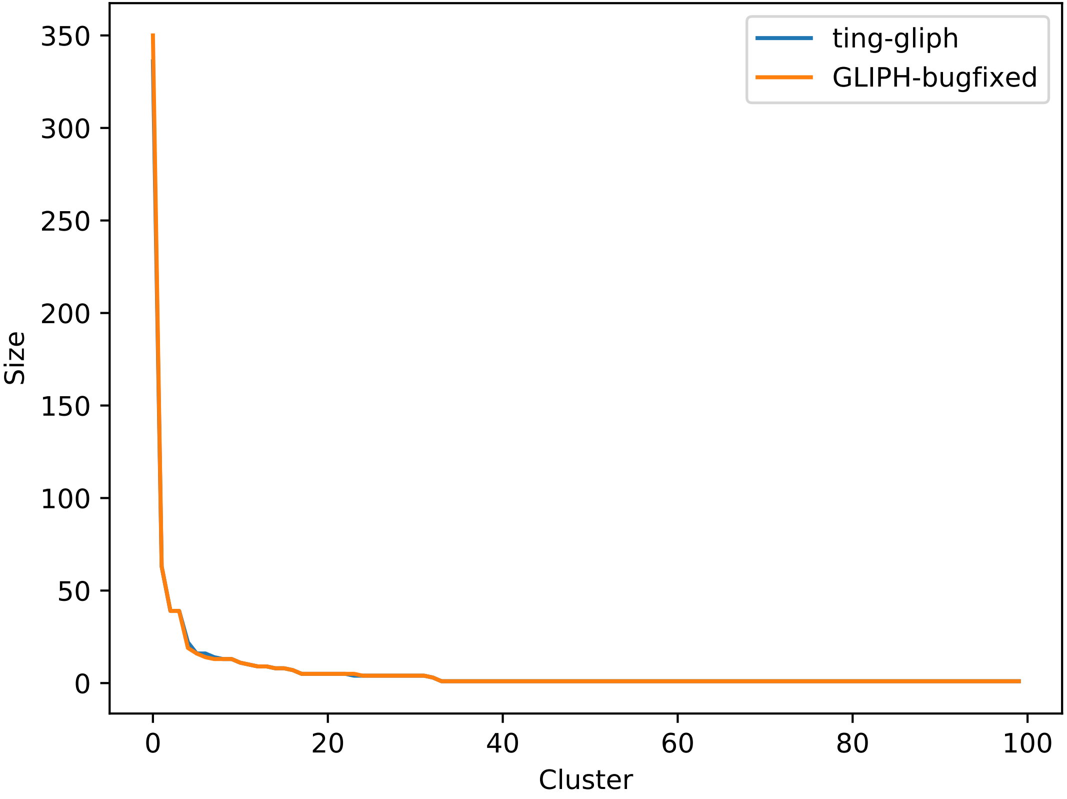

### specific_R205-L01-D733D533_cluster.pdf

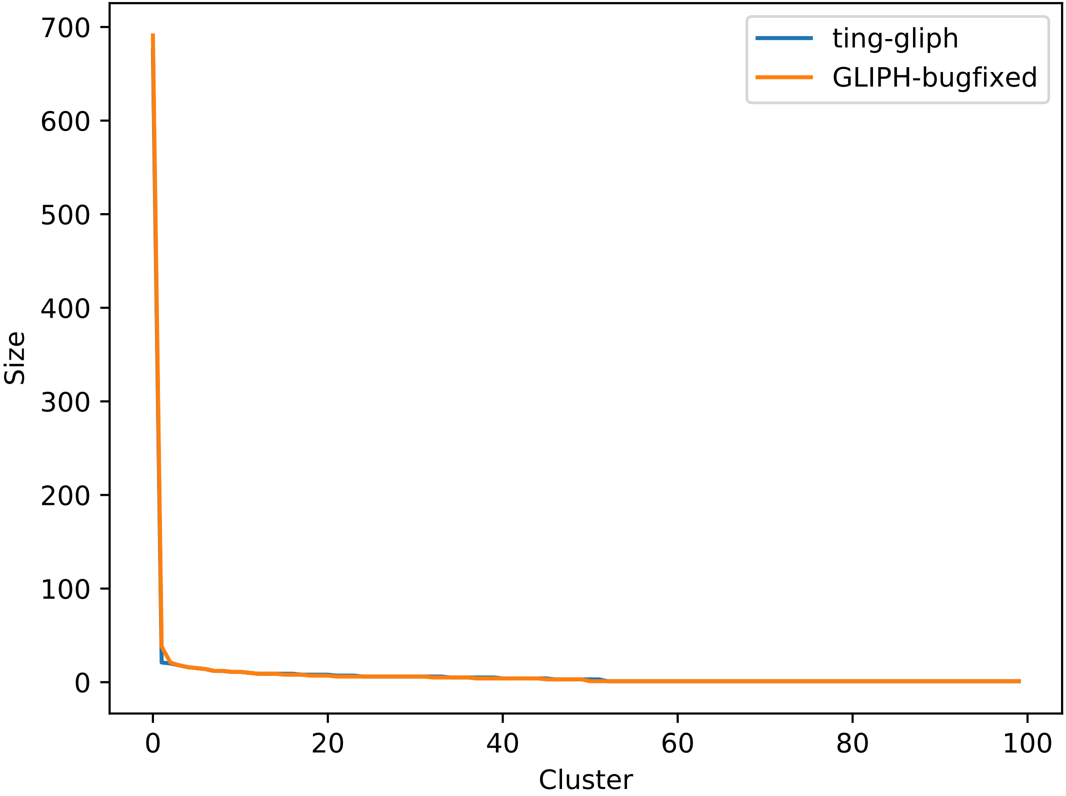

### specific_R205-L01-D781D581_cluster.pdf

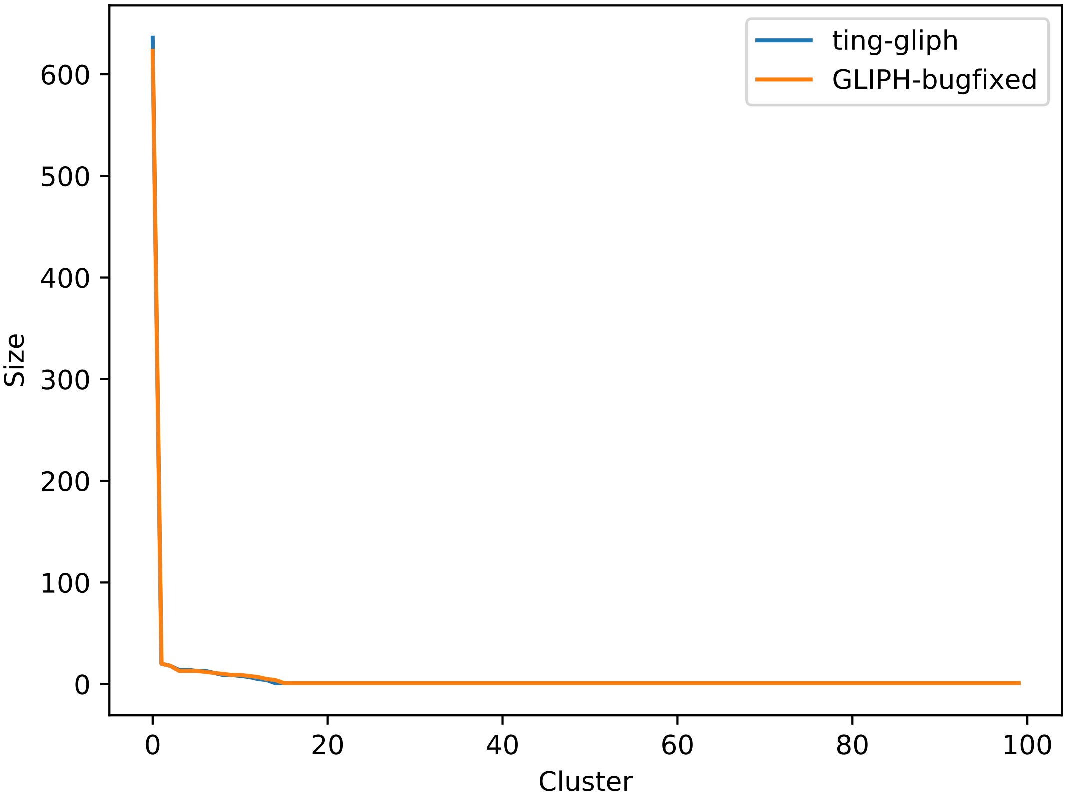

### specific_R205-L01-D783D583_cluster.pdf

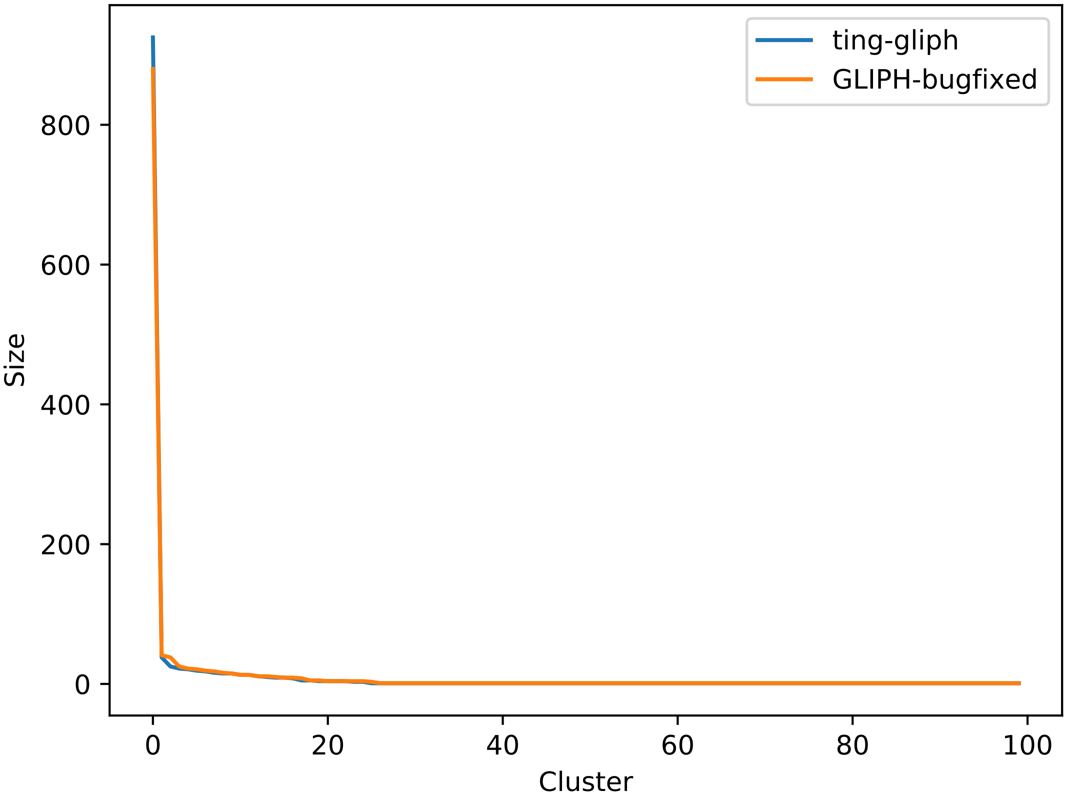

### specific_R205-L01-D785D585_cluster.pdf

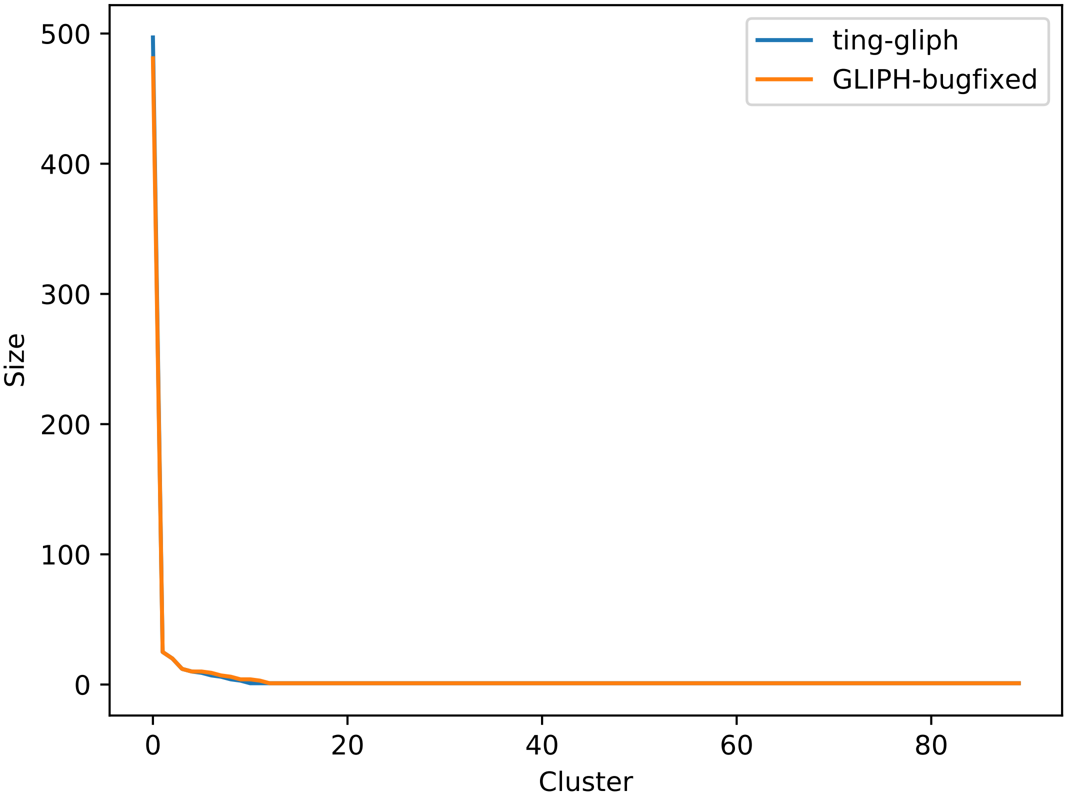

### specific_R206-L03-D701D501_cluster.pdf

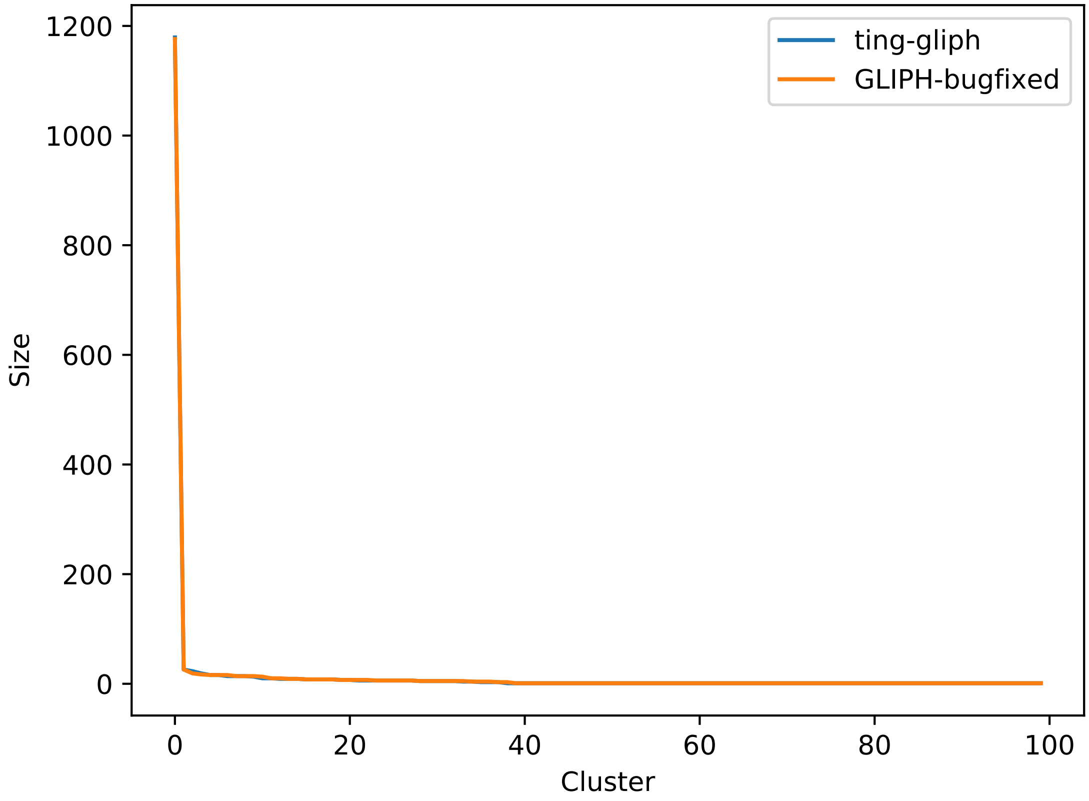

### specific_R206-L03-D721D521_cluster.pdf

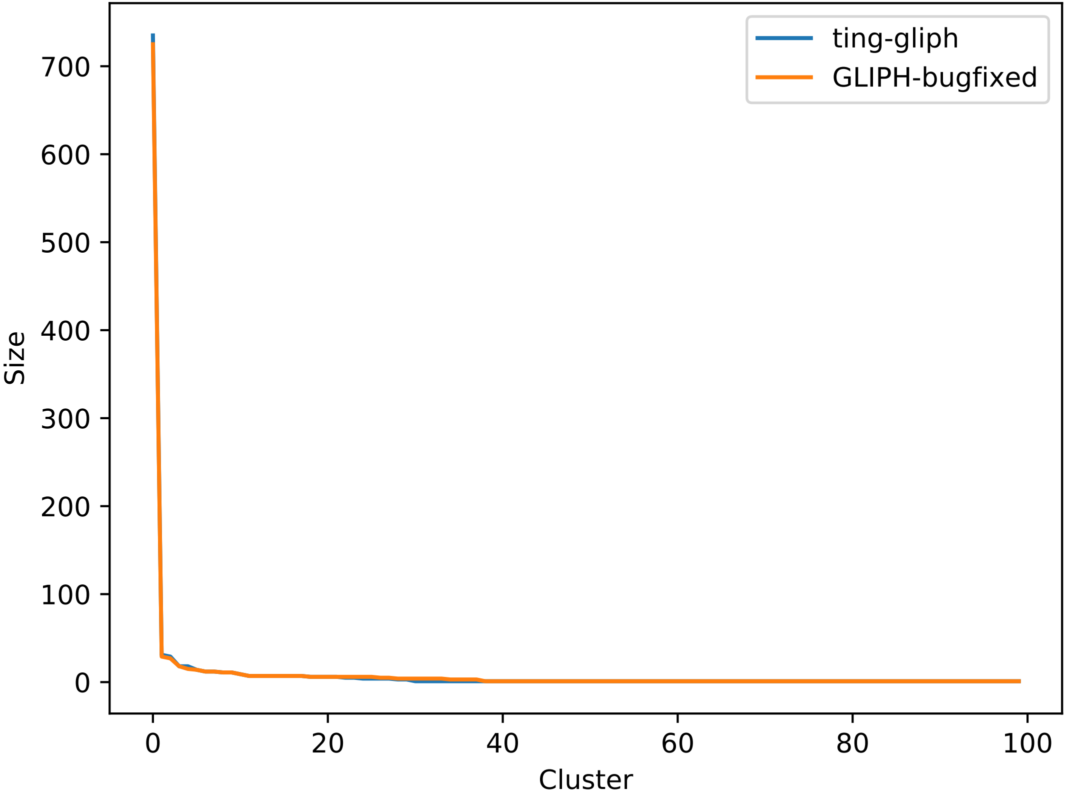

### specific_R206-L03-D725D525_cluster.pdf

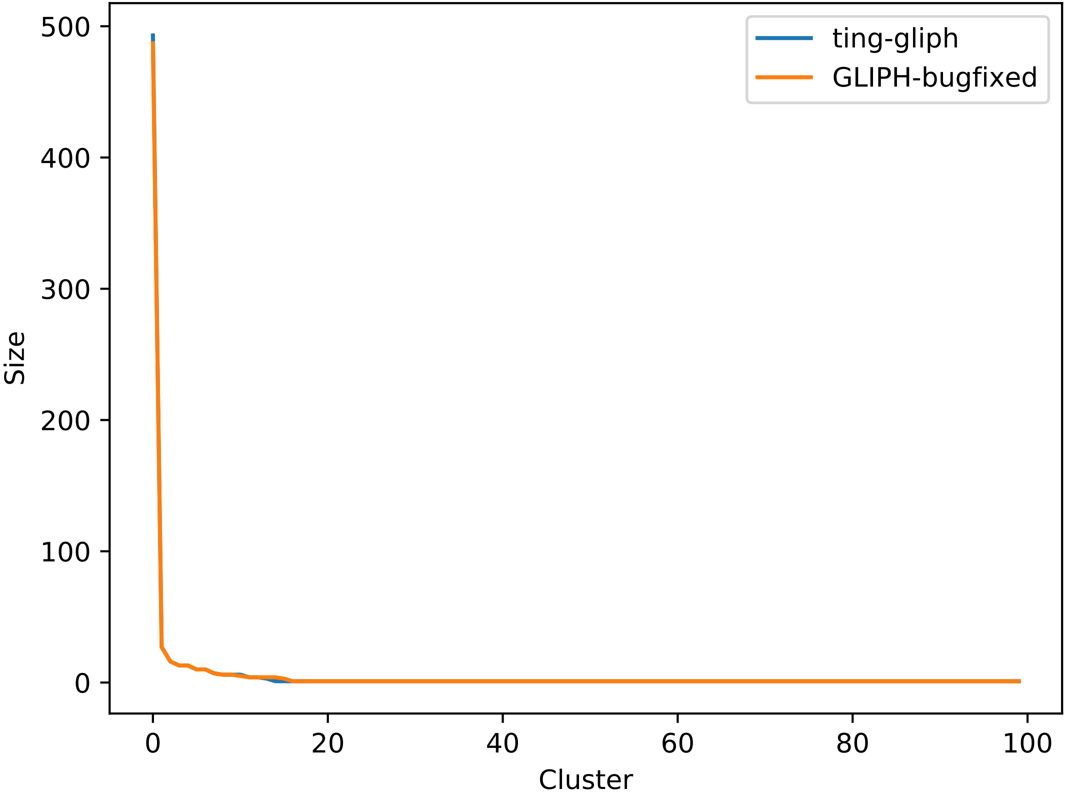

### unknown_LULO-TC-DN1_Tc_N_CD40L+_cluster.pdf

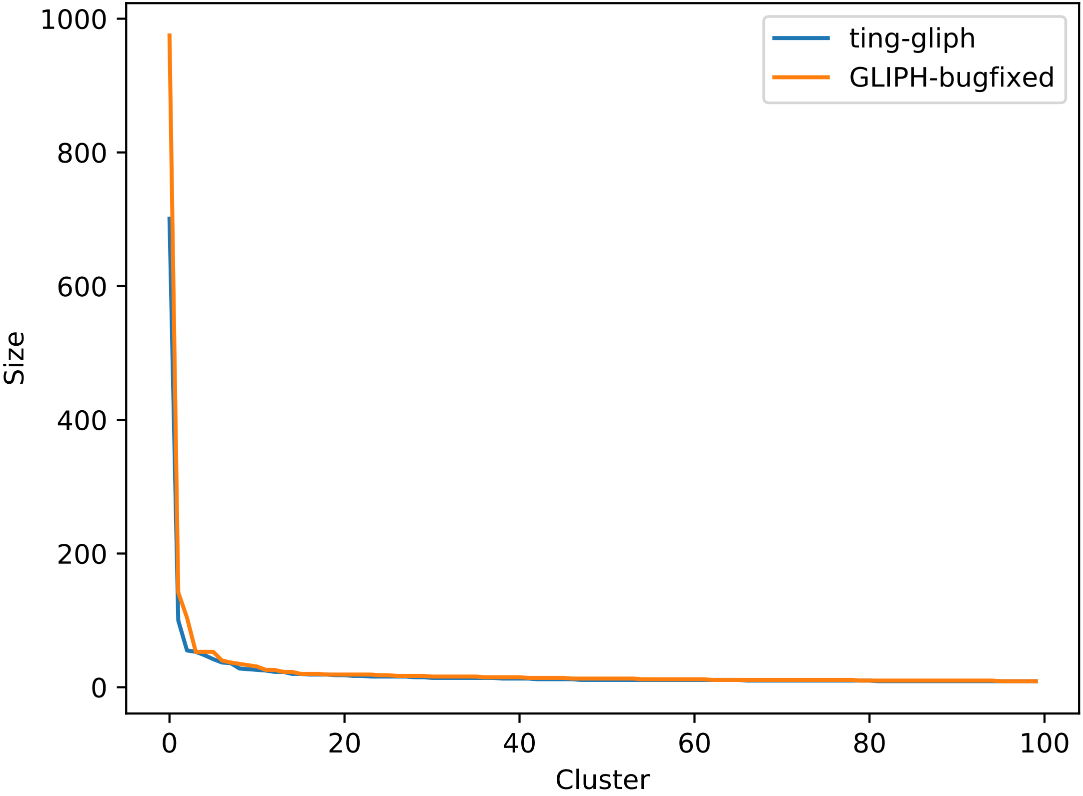

### unknown_LULO-TC-DN2_Tc_N_CD40L+_cluster.pdf

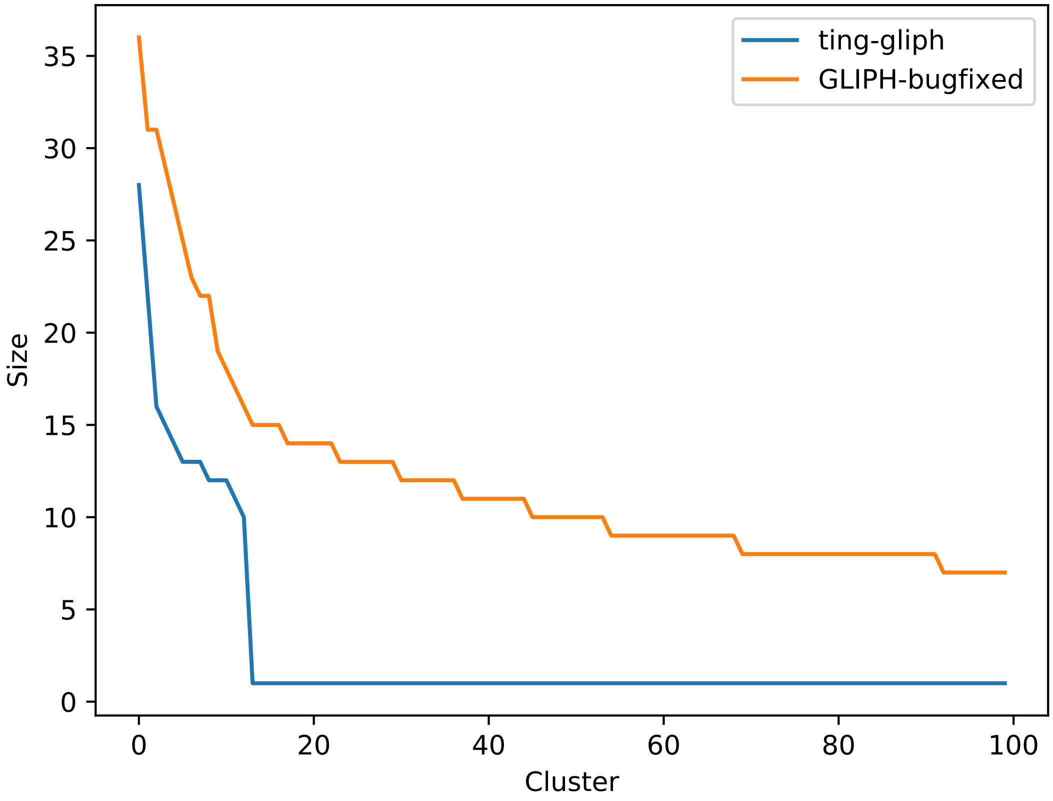

### unknown_LULO-TC-DN3_Tc_N_CD40L+_cluster.pdf

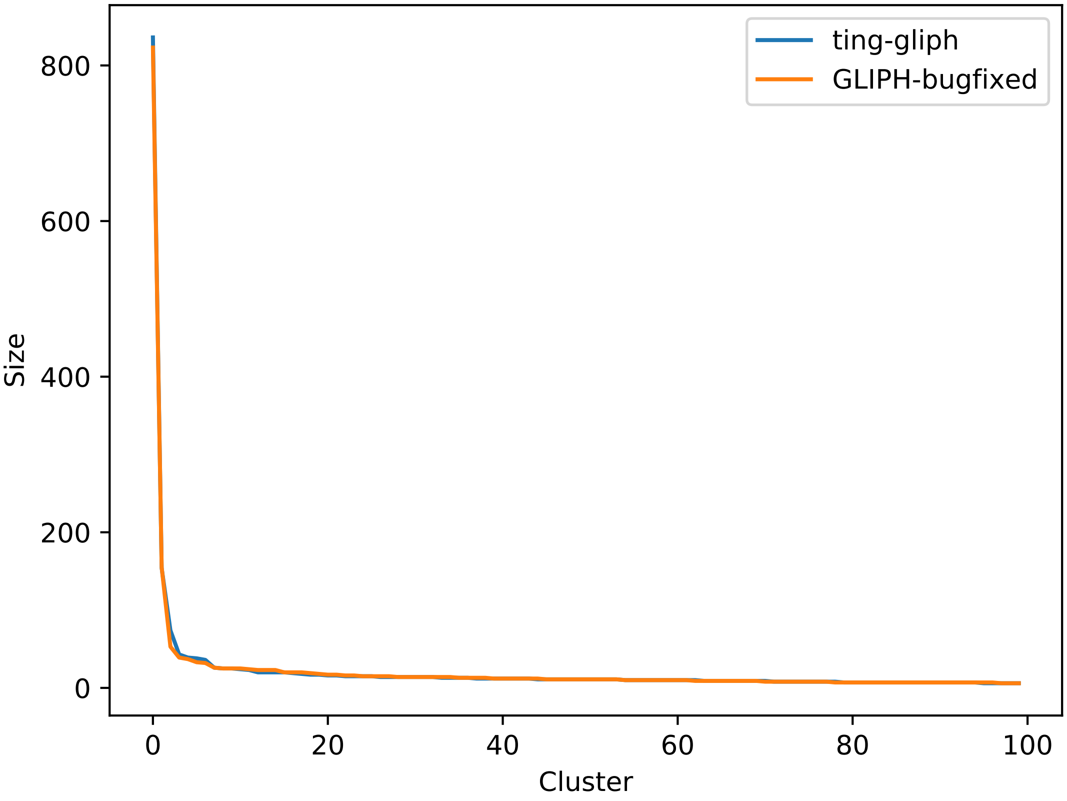

### unknown_LULO-TC-DN4_Tc_N_CD40L+_cluster.pdf

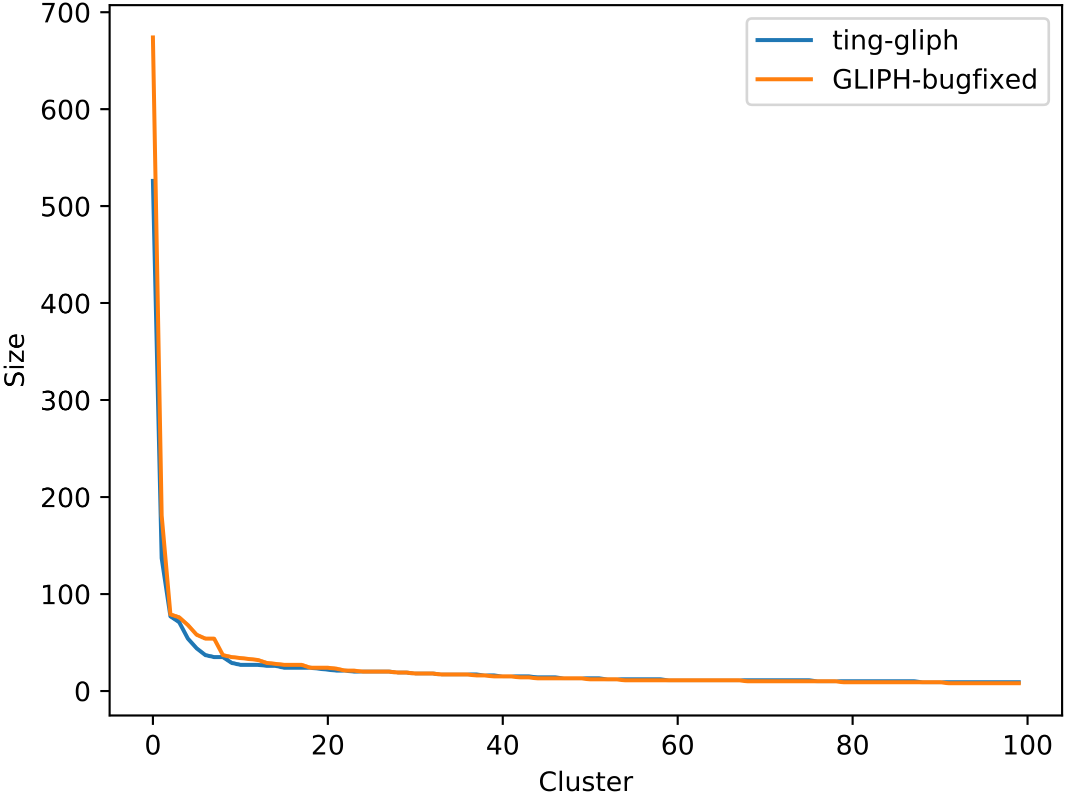

### unknown_LULO-TC-DN5_Tc_N_CD40L+_cluster.pdf

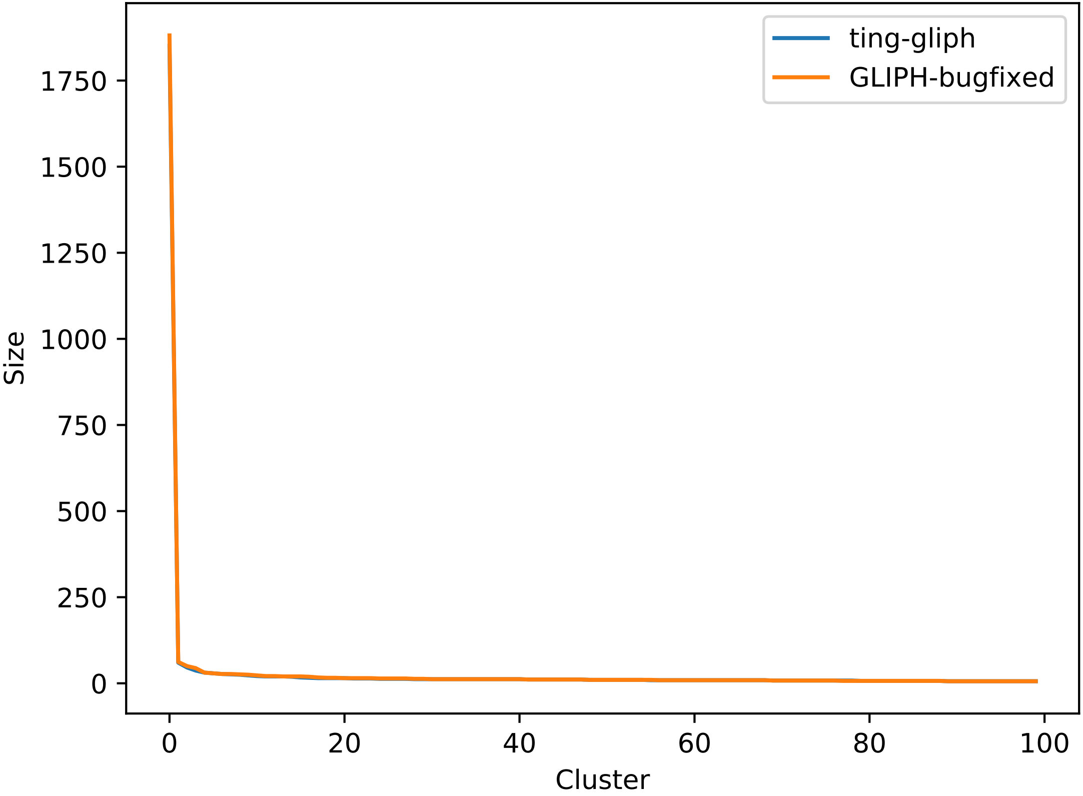

### unknown_LULO-TC-DN6_Tc_N_CD40L+_cluster.pdf

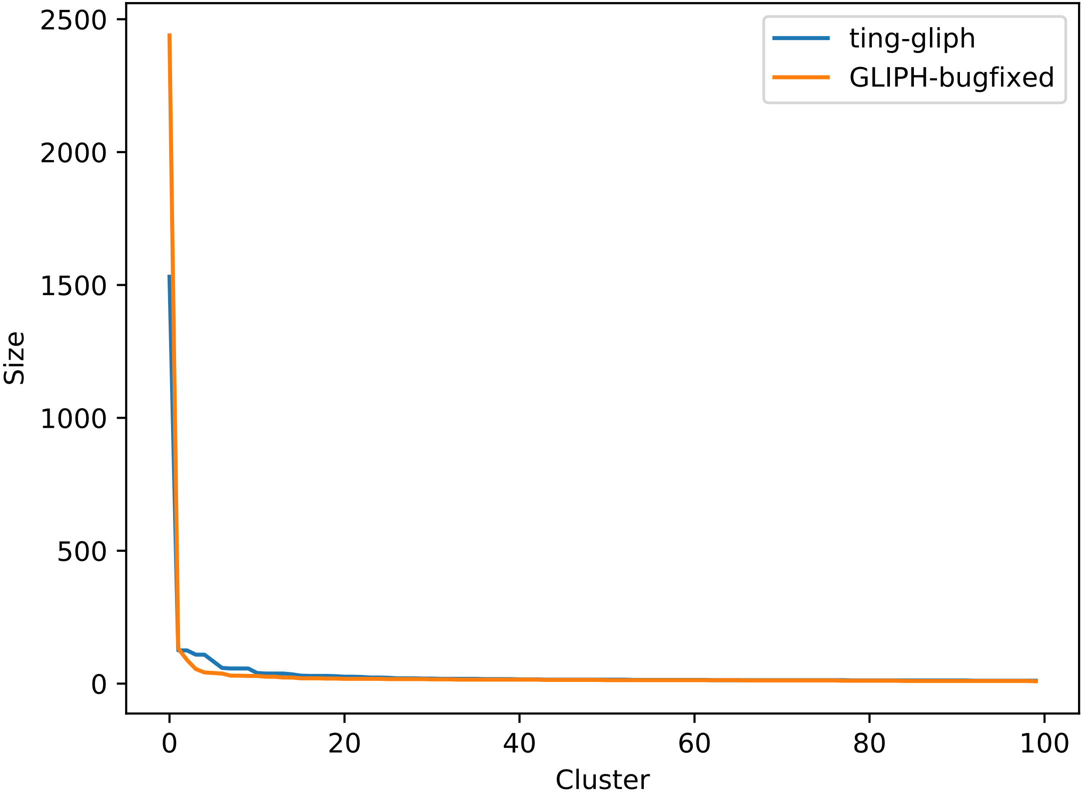
